# Quantification of phosphonate drugs by ^1^H-^31^P HSQC shows that rats are better models of primate drug exposure than mice

**DOI:** 10.1101/2022.01.30.478340

**Authors:** Yasaman Barekatain, Sunada Khadka, Kristen Harris, Jorge Delacerda, Victoria C. Yan, Ko-Chien Chen, Cong-Dat Pham, Md. Nasir Uddin, Rony Avritcher, Eugene J. Eisenberg, Raghu Kalluri, Steven W. Millward, Florian L. Muller

**Author notes:** Correspondence should be addressed to Florian L. Muller or Yasaman Barekatain.

## Abstract

The phosphonate group is a key pharmacophore in many anti-viral, anti-microbial, and anti-neoplastic drugs. Due to its high polarity and short retention time, detecting and quantifying such phosphonate-containing drugs with LC/MS-based methods is challenging and requires derivatization with hazardous reagents. Given the emerging importance of phosphonate-containing drugs, developing a practical, accessible, and safe method for their quantitation in pharmacokinetics (PK) studies is desirable. NMR-based methods are often employed in drug discovery but are seldom used for compound quantitation in PK studies. Here, we show that proton-phosphorous (^1^H-^31^P) heteronuclear single quantum correlation (HSQC) NMR allows for quantitation of the phosphonate-containing enolase inhibitor HEX in plasma and tissue at micromolar concentrations. Although mice were shown to rapidly clear HEX from circulation (over 95% in <1 hr), the plasma half-life of HEX was more than 1hr in rats and nonhuman primates. This slower clearance rate affords a significantly higher exposure of HEX in rat models compared to mouse models while maintaining a favorable safety profile. Similar results were observed for the phosphonate-containing antibiotic, fosfomycin. Our study demonstrates the applicability of the ^1^H-^31^P HSQC method to quantify phosphonate-containing drugs in complex biological samples and illustrates an important limitation of mice as preclinical model species for phosphonate-containing drugs.

## Introduction

The phosphonate moiety is a bioisostere of phosphate esters^1^, and phosphonate-containing drugs are emerging in a wide variety of health applications^2^, including antibiotics (fosfomycin^3^, fosmidomycin, SF2312^4^), anti-virals (tenofovir, adefovir, cidofovir nucleotide analogs^5^), and anti-neoplastic drugs^6^. Phosphate esters are found in many anabolic and catabolic intermediates making phosphonates an attractive replacement for these functionalities in drugs targeting aberrant metabolism. One example of such a drug is the small molecule enolase inhibitor HEX (**Figure 1a,b**), where the phosphonate binds to the Arg373 residue of the enolase enzyme. In preclinical studies, enolase 1 (*ENO1*)-deleted glioma tumors exhibited significant sensitivity to HEX^7,8^. Like many other phosphonate-containing drugs^3^, HEX has a favorable safety profile allowing high dose administration in rodent and primate models.

**Figure 1:**
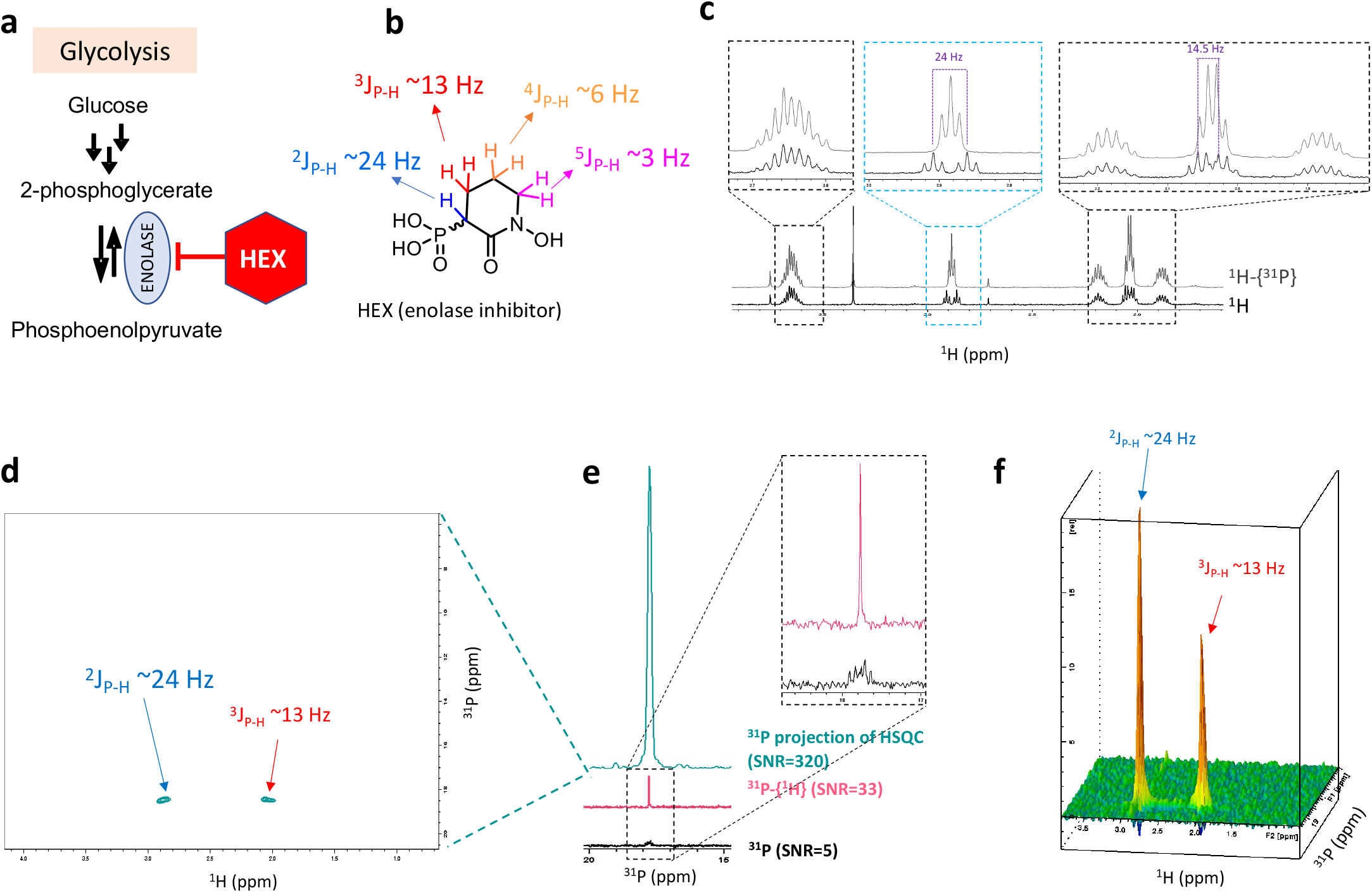
High SNR of the ^1^H-^31^P HSQC spectrum compared to 1D ^31^P spectra. (**a**) HEX selectively inhibits the glycolytic enolase enzyme. (**b**) The molecular structure of HEX with the J-coupling values between ^1^H and ^31^P nucleus. (**c**) proton (^1^H) and phosphorous-decoupled proton (^1^H-{^31^P}) spectra of HEX. (**d**) The ^1^H-^31^P HSQC spectrum of HEX where peaks associated with J2 and J3 coupling are evident. (**e**) The ^1^H-^31^P HSQC (acquisition time ~ 15 minutes) can achieve a significantly higher signal-to-noise ratio (SNR) compared to 1D ^31^P and proton-decoupled ^31^P (acquisition time ~ 20 minutes). (**f**) The oblique view of the ^1^H-^31^P HSQC spectrum in panel **d**. Higher intensity of the J2 peak of HEX can be seen.

Establishing the time-dependent plasma concentration of any drug is a crucial step in developing a pharmacokinetic model. However, the absolute quantification of electron-rich phosphonates is challenging by LC- and MS-based methods. Because of their high polarity and negative charge, these compounds have a low retention time in reverse phase HPLC columns leading to poor resolution from endogenous and abundant phosphate and phosphate esters in complex biological mixtures. While chemical derivatization of phosphonates to generate less polar compounds prior to analysis is commonly employed, these techniques require toxic chemicals such as trimethylsilyl diazomethane^7,9,10^. Although radioactive derivatives can be used to determine the PK of phosphonates in plasma and organs, these methods require dedicated radiosynthesis capability and instrumentation, which prohibits their routine use in most academic labs.

One-dimensional (1D) proton-decoupled phosphorus (^31^P-{^1^H}) NMR scan is commonly employed to study phosphonate-containing drugs and synthetic intermediates. However, the ^31^P nucleus suffers from very low sensitivity in ^31^P NMR, limiting the utility of this technique at low phosphonate concentrations^11^. Twodimensional (2D) NMR techniques such as heteronuclear single quantum correlation (HSQC) enhance detection of nuclei with lower NMR sensitivity (^13^C, ^15^N, and ^31^P) through polarization transfer of magnetization from the more sensitive ^1^H nucleus^12^. The ^1^H-^13^C HSQC scan has found utility in medicinal chemistry and cancer metabolism^13-15^, and ^1^H-^15^N HSQC is extensively used in protein structural studies^16^; however, ^1^H-^31^P HSQC has gained limited attention in biology and even in medicinal chemistry. The 2D ^1^H-^31^P NMR has been employed to study phosphate-ester endogenous metabolites^17-19^. In this technique, the ^3^J_H-P_ coupling constants are very small, limiting the efficiency of magnetization transfer, while the higher ^2^J_H-P_ coupling constants (~20 Hz) in phosphonate-containing drugs offer better magnetization transfer. Thus, ^1^H-^31^P HSQC for phosphonate-containing drugs combines the sensitivity of the ^1^H-nucleus with the specificity of ^31^P. We hypothesize that compared to the 1D proton-decoupled ^31^P, ^1^H-^31^P HSQC is more sensitive to detecting phosphonate compounds in biological samples.

Here, we illustrate the capability of ^1^H-^31^P HSQC to quantify micromolar levels of HEX and fosfomycin in plasma and tissues across different species (mouse, rat, nonhuman primate). Such studies were performed using a standard Bruker NMR system without pre-analysis chemical derivatization. This method allows for absolute quantification of the drug uptake in various organs. Our studies revealed that mouse models poorly predict the plasma clearance of HEX in primate models. In contrast, the rat models provide a very accurate estimate of compound clearance in primates, likely reflecting the actual human situation. The area under the curve (AUC24) for HEX was around five times higher in rats than in mice indicating higher drug exposure. Thus, our data suggest that the rats may provide a more suitable model to study the pharmacokinetics and pharmacodynamics of phosphonate drugs. This study is the proof-of-principle experiment to demonstrate the usefulness of the ^1^H-^31^P HSQC method and the importance of choosing the appropriate animal model to study the metabolism and clearance of phosphonate-containing drugs.

## Results and Discussion

### Maximum sensitivity for phosphonates achieved using an adapted ^1^H-^13^C HSQC sequence

First, we determined proton-phosphorous j-coupling constants (J_P-H_) of HEX by comparing the 1D proton (^1^H, black) and phosphorous-decoupled proton (^1^H-{^31^P}, gray) spectra (**Figure 1c**). From this spectrum, we obtained an approximate value of ^2^J_P-H_, which informed our estimate of the INEPT evolution time in 2D polarization transfer experiments. HEX has the ^2^J_P-H_ and ^3^J_P-H_ coupling constants of ~24 Hz and 13 Hz, respectively. Then, we used the ^1^H-^13^C HSQC pulse sequence to set up the ^1^H-^31^P HSQC, as explained in the method section. **Figure 1d** shows the ^1^H-^31^P HSQC spectrum of HEX acquired with the HSQCETGP pulse program, where the peaks associated with the ^2^J_P-H_ and ^3^J_P-H_ correlations are obvious and well-resolved. We then obtained the 1D ^31^P spectral projections from the ^1^H-^31^P HSQC spectrum (green spectrum in **Figure 1d**) and compared it with 1D ^31^P (black) and proton-decoupled ^31^P (^31^P-{^1^H}, pink) spectra. A 10-fold increase in sensitivity for the ^31^P nucleus was obtained in the HSQC method relative to the 1D ^31^P and ^31^P-{^1^H} experiments (**Figure 1e**). The higher intensity of the ^2^J_P-H_ peak can be better appreciated when the ^1^H-^31^P HSQC spectrum is shown in the oblique view (**Figure 1f**). This HSQC sequence is available on Bruker NMR spectrometers, allowing this method to be implemented at most academic centers.

### Determining *in vivo* pharmacokinetics of HEX using ^1^H-^31^P HSQC

Given the high sensitivity of the ^1^H-^31^P HSQC method, we sought to use this method to evaluate the in vivo pharmacokinetics of HEX. Unlike the 1D proton spectrum, HSQC peak intensity cannot be directly used for quantification. Thus we sought to obtain the calibration curve which can be used to calculate the unknown concentrations of HEX in plasma. The ^31^P spectral projections of known concentration of HEX (**Supplementary Figure S1**) were used to determine the dose calibration curve in **Figure 2a,** with the x-axis indicating the integral of the HEX peak from the ^31^P spectral projection of the ^1^H-^31^P HSQC spectrum and the corresponding known HEX concentration on the y-axis. The lower detection limit was determined to be about 25 μM HEX (for 200 μL plasma, HSQCETGP pulse program, ns=128, and regular 5 mm NMR tube), and a minimum signal to noise ratio (SNR) of 5 was considered for a reliable signal. To obtain the PK profile of HEX in NHP, the *Macaca fascicularis* monkey was injected with 100 mg per kg HEX subcutaneously. After injection, blood was collected at different time points, and hydrophilic extracts were prepared for the NMR studies (see methods). The NMR studies can also be done on intact plasma, albeit with a reduction in the signal-to-noise in the ^31^P resonance (**Supplementary Figure S2**). The proton spectrum of plasma extract is illustrated in **Figure 2b,** where HEX peaks are obscured by highly abundant metabolites, hindering the detection and quantification of HEX in plasma. **Figure 2c** shows the ^1^H-^31^P HSQC spectrum of the same sample where peaks from ^2^J_P-H_ and ^3^J_P-H_ of HEX and endogenous phosphodiester (PE) are detected. Using the ^1^H-^31^P HSQC experiment, 1D ^31^P spectral projections were obtained for various time points post-injection (**Figure 2d**). As expected, the intensity of the HEX peak in the plasma decreased over time following the last dose. Using the calibration curve, we calculated plasma concentrations of HEX for each time point (**Figure 2e)**.

**Figure 2:**
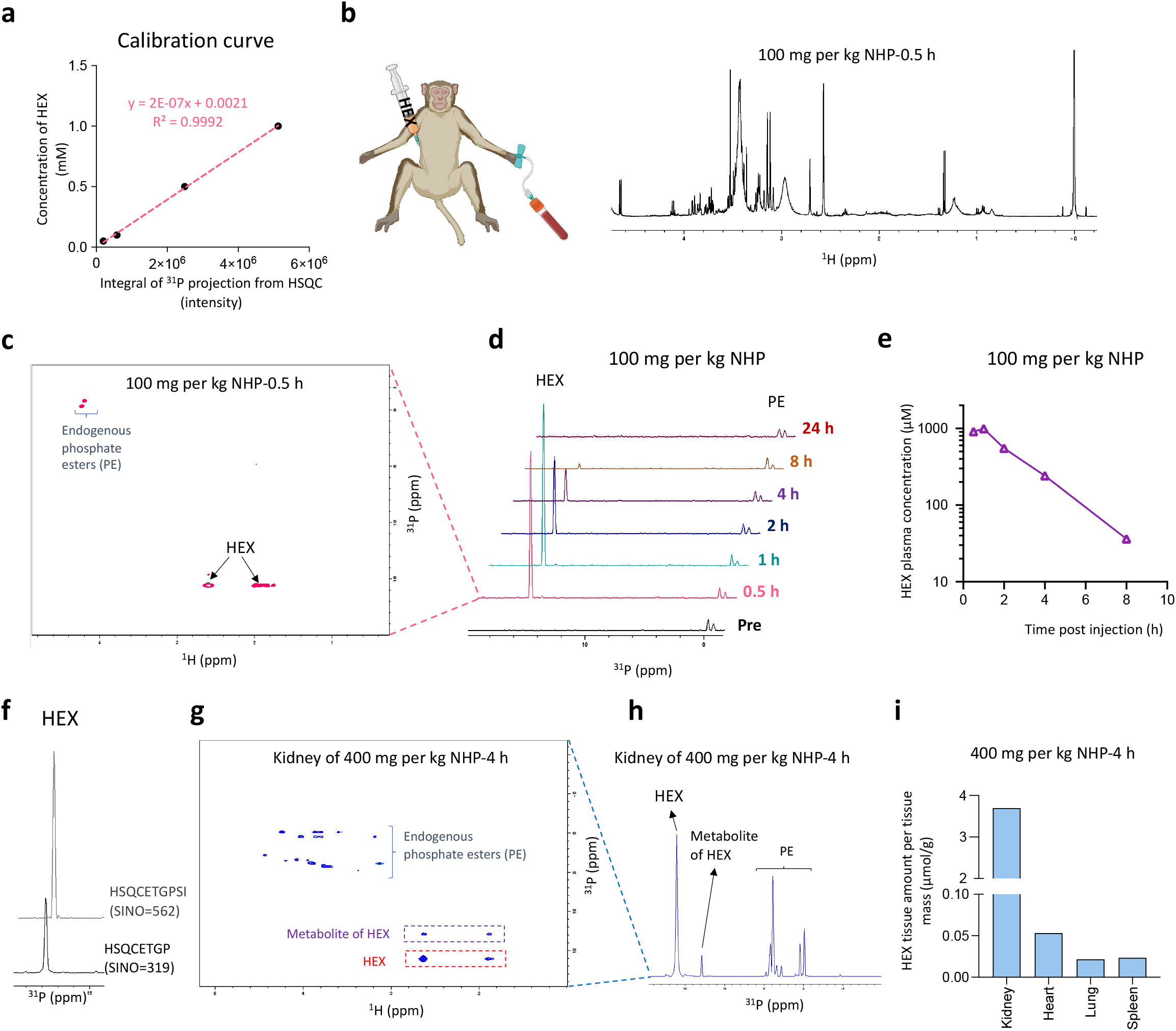
Determining *in vivo* pharmacology of HEX using ^1^H-^31^P HSQC. (**a**) Dose calibration curve used to calculate concentration of HEX in plasma. (**b-e**) A male cynomolgus monkey (NHP) was subcutaneously injected with 100 mg per kg, with plasma collected at the indicated times. (**b**) Even at the earliest time points, HEX peaks are unidentifiable in the proton spectrum of a plasma extract due to the presence of highly abundant metabolites. (**c**) In contrast, the ^1^H-^31^P HSQC spectrum of the same sample is highly specific, detecting only phosphorous-containing compounds in plasma. (**d**) The ^31^P intensity projection of ^1^H-^31^P HSQC spectra was used to calculate HEX concentration. (**e**) Plasma concentration of HEX in an NHP at various time points after the final dose. (**g**) The HSQCETGPSI pulse sequence yields higher SNR. (**h**) The ^1^H-^31^P HSQC spectrum of a NHP’s kidney, 4 hours post-injection with 400 mg per kg HEX. (**i**) The positive projection of all columns from the ^1^H-^31^P HSQC spectrum where HEX and its metabolite are shown. (**j**) HEX levels across various tissues from an NHP subcutaneously injected with 400 mg per kg HEX.

To further increase the detection sensitivity of HEX in biological samples, we carried out the ^1^H-^31^P HSQC experiment using the ^1^H-^13^C HSQCETGPSI pulse sequence (where SI stands for sensitivity increased, **Figure 2f**). This modified experiment was used to measure HEX concentrations in ex vivo tissues taken from an NHP injected with 400 mg per kg HEX (**Figure 2g-I** and **Supplementary Figure S3**) and sacrificed 4hrs later. Here, we illustrate the HSQC technique’s capability to measure the micromolar concentration of phosphonate-containing drugs in plasma and tissues. While such measurements are possible using LC/MS-based methods, our results show that this can be done with a standard NMR.

### HEX has a more favorable, human-like PK profile in rats and NHPs compared to mice

Having established the efficient method to detect HEX and its PK profile in biological samples, we sought to compare the pharmacology of HEX across different species to determine the optimum animal model for PK studies. The single dose of 150 mg per kg HEX was injected subcutaneously in mice, rats, and NHP, and plasma was collected at various time points. We then extracted the plasma and prepared the samples for the NMR studies (**Figure 3a-c**). Consistent with findings from **Figure 1i** (an NHP injected with 100 mg per kg HEX), the plasma elimination half-life of HEX was determined to be 77 minutes in NHP. Surprisingly, we found that HEX has a plasma half-life in rats (79 min) comparable to NHP. Finally, HEX has the least favorable metabolism in the mouse model, with a half-life as short as 22 minutes. **Figure 3d** compares the HEX plasma concentration in NHP, rat, and mouse models injected with a single dose of 150 mg per kg HEX alongside the estimated PK profile of HEX in humans. The *in vitro* IC_50_ values for cancer cell lines homozygous *ENO1*-deleted (*ENO1*-/-), heterozygous *ENO1*-deleted (*ENO1*+/-), and *ENO1*-wildtype (*ENO1*+/+) are indicated as the red, gray, and black dashed lines in **Figure 3d**. Roughly, this suggests the dose required for therapeutic efficacy against these phenotypes and shows that in NHP, therapeutic plasma concentrations are maintained in plasma for 12-24 hours at no-observed-adverse-effect level (NOAEL) dosing levels.

**Figure 3:**
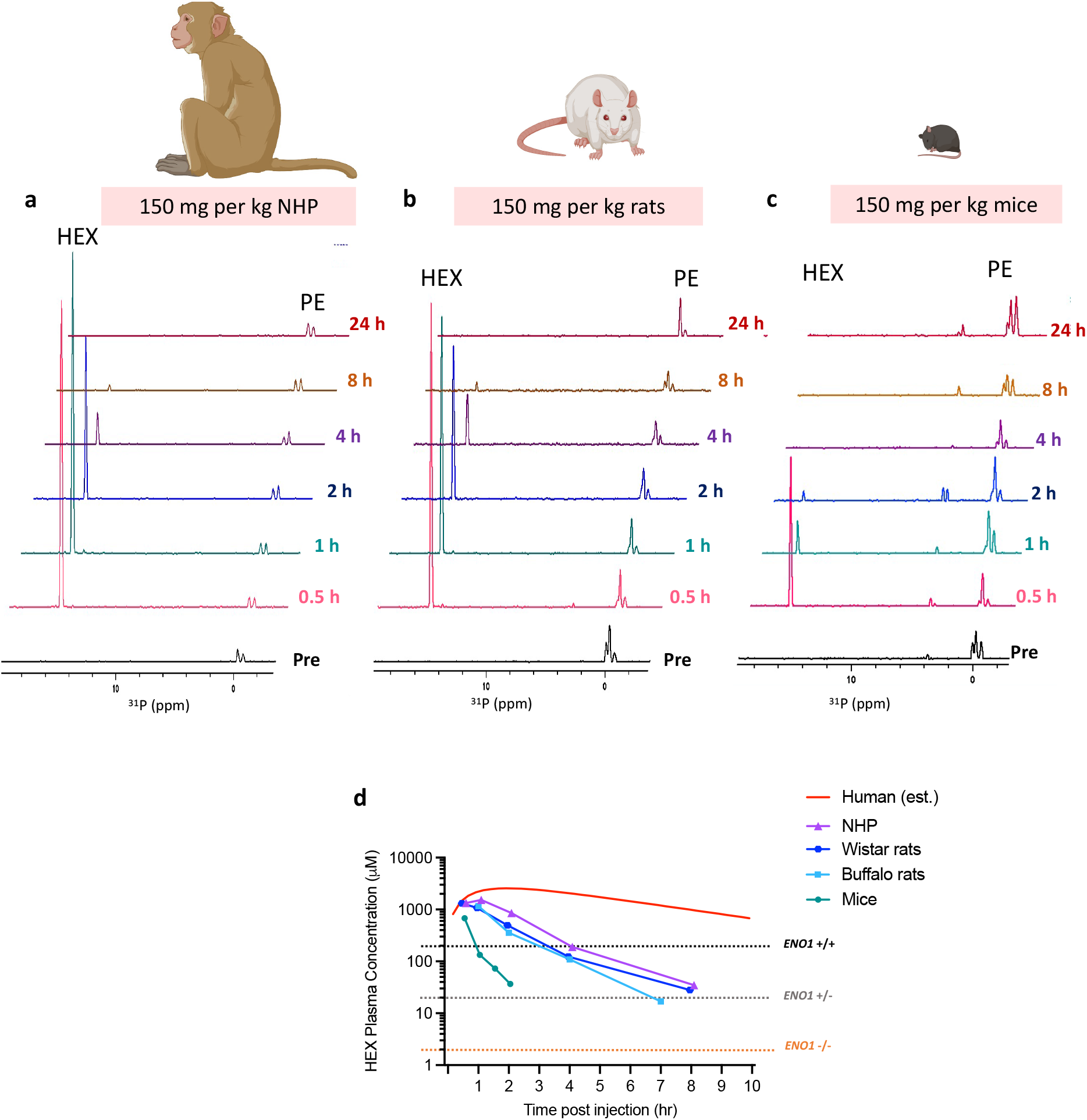
More favorable PK profile of HEX in NHPs and rats versus mice. Monkey, rats, and mice were injected with 150 mg per kg of HEX subcutaneously. Plasma was collected at various time points post-injection and prepared for NMR studies to find HEX plasma concentration and PK profile. (**a-c**) The positive projection of all columns of ^1^H-^31^P HSQC spectra of plasma extract was collected from (**a**) a cynomolgus monkey, (**b**) Wistar rats, and (**c**) black mice. HEX has a similar drug elimination rate in the plasma of an NHP and rats. (**d**) Comparison of PK profile of HEX in a monkey, rats, mice, and humans (estimated) based on single 150 mg per kg subcutaneous dose. Estimated human data were calculated using the one-compartment PK model. The dashed lines indicate the cell-based IC50 in homozygous *ENO1*-deleted (*ENO1*-/-), heterozygous *ENO1*-deleted (*ENO1*-/+), and *ENO1*-wildtype (*ENO1*+/+) cancer cells (from Lin et al, Nature Metabolism 2020).

Moreover, using the predicted values in **Supplementary Table S1** and **S2**, we estimated the multidose HEX PK at the maximum recommended starting dose in humans shown in **Supplementary Figure S4**. Pharmacokinetic properties of HEX among NHP, rats, and mice are compared in **Table1**. Our findings demonstrate that HEX has a similar PK profile in rats and an NHP with a longer drug elimination half-life and higher exposure than mice. While the tumor model studies are not currently possible in NHP, these studies suggest that rat models may better predict HEX efficacy.

**Table 1:** Pharmacokinetic properties of HEX among different pre-clinical species. The non-compartmental analysis was used to determine clearance and volume of distribution (CL and VD).

|  |  |  |  |
| --- | --- | --- | --- |
| Species | Mice | Rat* | NHP |
| Order | Rodentia | Rodentia | Primate |
| Strain | C57/BL | Wistar<br>Buffalo | Non-native<br>Cynomolgus |
| Gender | Male | Male | Male |
| Weight<br>(kg) | 0.025-0.030 | 0.300-0.325 (Wistar)<br>0.54-0.56 (Buffalo) | 2.87 |
| S.C. HEX NOEFL<br>Tolerability (mg/kg)** | 450 | >150 | 200 |
| Kel | 1.87 | 0.53 | 0.54 |
| Plasma elimination<br>half-live (min)*** | 22 | 79 | 77 |
| Plasma C<sub>max</sub><br>(μM) | 676 | 1317 | 1337 |
| Plasma C<sub>min</sub><br>(μM) | 37 | 24 | 35 |
| AUC<sub>(0-t)</sub><br>(μM.h) | 487 | 2568 | 4021 |
| AUC<sub>(0-inf)</sub><br>(μM.h) | 506 | 2615 | 4085 |
| AUMC<sub>(0-t)</sub><br>(μM.h) | 381 | 4327 | 9108 |
| AUMC<sub>(0-inf)</sub><br>(μM.h) | 421 | 4726 | 9647 |
| MRTINF<br>(h) | 0.83 | 1.81 | 2.36 |
| CL<br>(L/h/kg) | 1.55 | 0.3 | 0.19 |
| VD<br>(L/kg) | 1.29 | 0.54 | 0.45 |

\* Data from male rats were pooled \*\* Single dose subcutaneous dose \*\*\* Calculated based on single 150 mg per kg subcutaneous dose

### Applicability of the ^1^H-^31^P HSQC method to detect other phosphonate-containing drugs

The phosphonate moiety is an essential functionality in a growing number of drugs. **Figure 4** shows the ^1^H-^31^P HSQC spectra of three common phosphonate-containing drugs with different chemical structures: zoledronic acid (isopropyl inhibitor for the treatment of osteoporosis and bone malignancies), fosfomycin (antibiotics), and tenofovir (nucleotide analog anti-viral for the treatment of hepatitis B). We first determined the proton and phosphorus J-coupling by comparing the proton and ^31^P-decoupled proton spectra. Then we performed ^1^H-^31^P HSQC using appropriate acquisition parameters for each compound. This figure illustrates that the 1D ^31^P spectrum projected from HSQC has a much higher SNR than 1D ^31^P and ^31^P-{^1^H}. These data indicate that the ^1^H-^31^P HSQC method is a general solution to the challenges associated with phosphonate drug quantitation.

**Figure 4:**
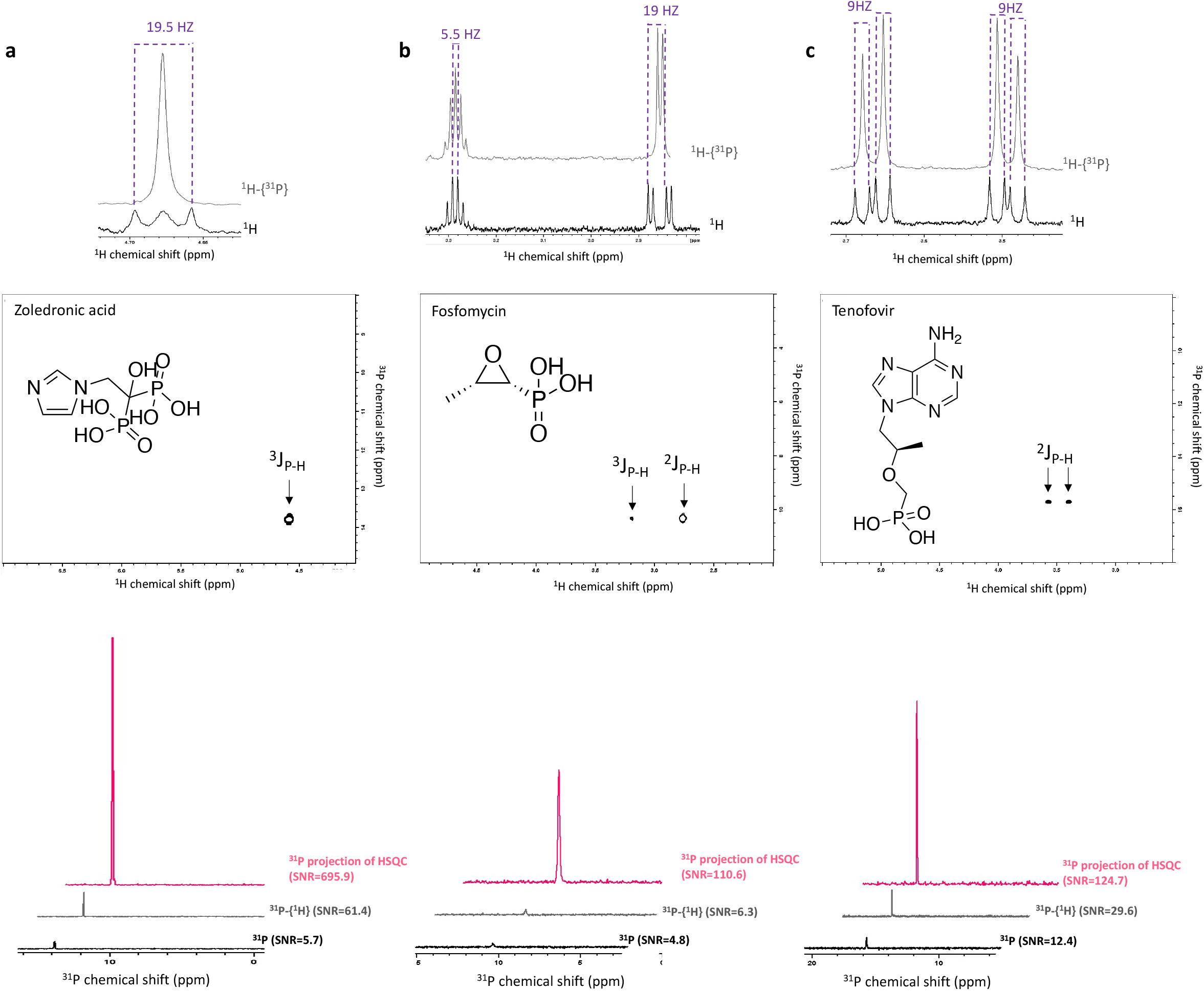
^1^H-^31^P HSQC method is applicable to detect and quantify other phosphonate-containing drugs besides HEX. Proton (^1^H), phosphorus-decoupled proton (^1^H-{^31^P}), and ^1^H-^31^P HSQC spectra of (**a**) zoledronic acid (≈ 400 μM), (**b**) Fosfomycin (≈ 350 μM), and (**c**) tenofovir (≈ 400 μM). The significantly higher SNR achieved from ^1^H-^31^P HSQC (HSQCETGPSI) spectra compared to 1D ^31^P and proton-decoupled phosphorous (^31^P-{^1^H}) spectra.

We next sought to use the method to determine the PK profile of fosfomycin (**Figure 5a**) in mice and rats injected subcutaneously with a single dose of 300 mg per kg (**Figure 5b-d**). Using the calibration curve shown in **Figure 5e**, we calculated the amount of fosfomycin in plasma at various time points after a single injection (**Figure 5f**). As expected, we found that fosfomycin has a slower plasma clearance rate in rats than mice, corroborating our findings with HEX. This figure further emphasizes that rats could be a better preclinical model to study the efficacy of phosphonate-containing drugs.

**Figure 5:**
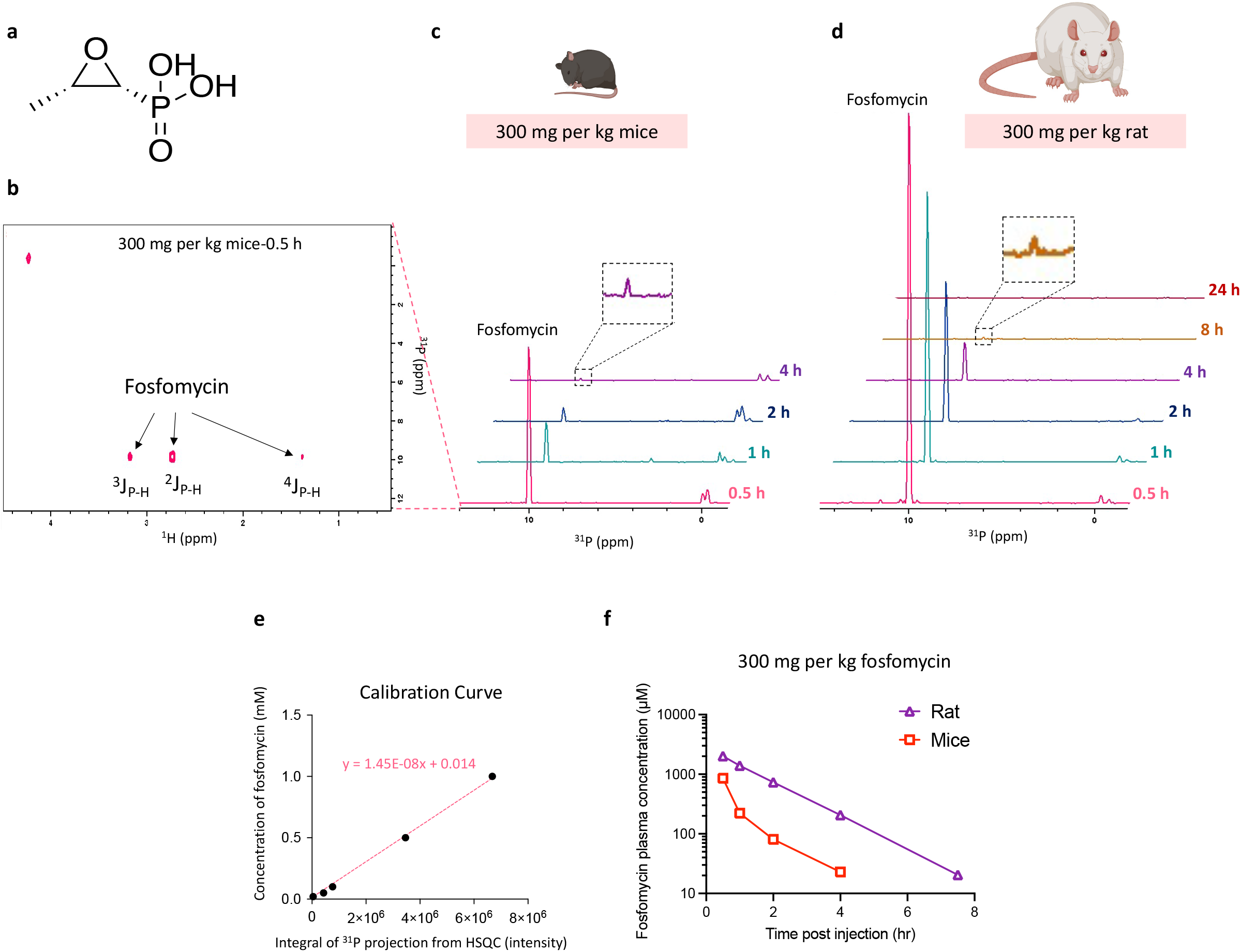
Fosfomycin has a more favorable PK profile in rats compared to mice, echoing the situation with HEX. (**a**) The molecular structure of fosfomycin. (**b**) The ^1^H-^31^P HSQC spectrum of mouse plasma 30 minutes after subcutaneous injection with a single dose of 300 mg per kg fosfomycin. J2, J3, and J4 coupling can be seen in the spectrum. (**c** and **d**) The positive projection of all columns of ^1^H-^31^P HSQC spectra of plasma extract in male and female black mice and a male buffalo rat injected with 300 mg per kg fosfomycin. (**e**) Calibration curve used to calculate unknown concentration of fosfomycin in plasma. (**f**) Fosfomycin plasma concentration in a rat and mice.

## Conclusion

Our study describes a new NMR-based method to quantify phosphonate-containing drugs in plasma and tissues using a standard NMR spectrometer and straightforward sample preparation. Using this method, we measured the PK profile of HEX in various species. We found that, unlike mice, rats slowly clear the drug from circulation and have a similar PK profile as NHPs. Our findings suggest that the short drug exposure of HEX in mice likely underestimates the efficacy of HEX in anti-neoplastic preclinical efficacy studies. Our studies also indicate that this may be true for other phosphonate drugs such as fosmidomycin and fosfomycin. Future studies will focus on using tumor-bearing rat models to test the efficacy of HEX in *ENO1*-deleted, *ENO1* heterozygous, and *ENO1* wild-type tumors. This method is also useful to study the enzyme activity of phosphonate drugs^20,21^. Moreover, the method described in this paper can be applied *in vivo* to study the drug distribution. Because our method provides higher sensitivity and more structural information, it is also suitable for detecting metabolites, provided that they retain the phosphorus atom. The method’s sensitivity can be significantly increased using an NMR spectrometer with a higher magnetic field and equipped with a 3 mm cryoprobe and Shigemi tubes and extracting more samples.

## Methods

### Experimental animals

Experimental procedures on the following animals were performed through a fee-for-service by Charles River Laboratories with approval of Charles River’s Institutional Animal Care and Use Committee (IACUC): male non-naïve cynomolgus monkeys (*Macaca fascicularis*) weighting 2.5-3.5 kg, male Wistar rats weighing 300-325 g, and male C57/BL mice weighting 25-30 g. Animal experiments using nude *Foxn1* BALB/c female mice, C57/BL mice, and male Buffalo rats weighing 500-550 g were performed at the University of Texas MD Anderson Cancer Center with MD Andersons’s approval IACUC.

### Administration and PK determination of phosphonate drugs

The 150 mg per mL HEX stock solution in water was prepared and sent to Charles Rivers. The stock solution was further diluted in saline to the desired concentration with PH of 7.2-7.4, followed by filtration using a 0.22 μm filter. Doses of 150 mg per kg per day or 100 mg per kg per day were injected in study animals (male C57/BL mice, male non-naïve cynomolgus monkeys, and male Wistar and buffalo rats). The 150 mg per mL fosfomycin stock solution was prepared in water with PH 7.2-7.4. A dose of 300 mg per kg was administrated in a mix of male and female C57/BL mice and a male buffalo rat. Blood samples were collected before injection and up to 24 hours post-injection. Plasma was separated from blood, snap-frozen, and kept at −80 °C for future use.

### Hydrophilic extract preparation of plasma

We adapted the hydropathic extraction method of BIDMC’s metabolomics core to prepare samples for HSQC analysis. Snap-frozen plasma thawed at room temperature. 200 μl of plasma placed in the new Eppendorf tube, and extracted using 400 μl of −20 °C pre-cold methanol, vortexed, and incubated in −20 °C for 45 min. The mixture was then vortexed and centrifuged at maximum speed (17,000*g*) for 30 min at 4°C. The clear supernatant was separated and dried in the vacuum concentrator (Eppendorf Vacufuge Plus). For NMR studies, the dried supernatant was dissolved in 470 μL D2O (Sigma Aldrich,151882) and 30 μL D2O with 3%TPS (Sigma Aldrich, 450510), and transferred in a standard 5 mm NMR tube (Wilmad-LabGlass, 16-800-497).

### Tissue extraction

snap-frozen tissues and tumors were extracted using 4 ml of dry-ice cold 66% methanol (66% methanol/34% water) for 200 mg of tissue. Tissues were then homogenized while kept in dry ice to avoid heating the samples. The homogenized mixture was then vortexed and kept at −80 °C for 24 hours. We then centrifuged the mixture at maximum speed (17,000*g*) for 1 hour at 4°C. The supernatant was then separated and dried using the vacuum concentrator (Eppendorf Vacufuge Plus). For NMR studies, the dried supernatant was dissolved in 470 μL D2O (Sigma Aldrich,151882) and 30 μL D2O with 3%TPS (Sigma Aldrich, 450510) and transferred in a standard 5 mm NMR tube (Wilmad-LabGlass, 16-800-497).

### NMR spectral acquisition

NMR studies performed on a 500 MHz Bruker Avance III HD spectrometer equipped with cryoprobe broadband observe probe located at the University of Texas MD Anderson Cancer Center. The 1D proton (^1^H) spectrum acquired zg30 pulse program, 1D phosphorous-decoupled (inverse gated) proton (^1^H-{^31^P}) spectrum obtained with zgig pulse program. 1D phosphorous (^31^P) and 1D proton-decoupled phosphorous (^31^P-{^1^H}) spectra are acquired using zg30 and zgpg30 pulse sequences. The 2D ^1^H-^31^P HSQC was adapted using ^1^H-^13^C HSQC (HSQCETGP and HSQCETGPSI where SI stands for sensitivity increased) pulse sequences. To acquire ^1^H-^31^P HSQC, we first change the nucleus from ^13^C to ^31^P, then set following parameters ns = 128 scans, gpz2 % = 32.40 (calculated using the au program “gradratio”), ^31^P SW = 40 ppm, O2p = 0 ppm, cnst2 = 22.95 which is adjusted for HEX ^2^J_P-H_ coupling, followed by “getprosal” commend). After the acquisition, we obtained the one-dimensional ^31^P spectral projection from the ^1^H-^31^P HSQC spectrum using “proj” commend in TopSpin, followed by baseline correction (abs). Integrals of HEX peak were used to calculate the concentration. NMR spectra were analyzed using TopSpin 3.1.

### Determining calibration curve

Two hundred micromolar of nontreated NHP plasma was spiked with the known concentration of HEX (1 mM, 0.5 mM, 0.1 mM, and 50 μM). It was then extracted using 400 μl methanol, followed by 45 minutes incubation at −20 ºC, and prepared for NMR studies. The 1H-31P HSQC spectrum was acquired for each sample using HSQCETGP and HSQCETGPSI pulse sequences with ns=128. From the ^1^H-^31^P HSQC spectrum, the 1D ^31^P spectral projections were obtained. The HEX peak in the 1D ^31^P projected spectrum was integrated and plotted versus the corresponding concentration of HEX. The line equation was obtained and used to calculate the unknown concentration of HEX in plasma and tissues.

### Prediction methodologies

The values for clearance (CL) and volume distribution (VD) in mice, rats, and NHPs were determined using the non-compartmental analysis. This analysis was chosen due to the limited number of data points. Below formulas were used to calculate CL and VD values:

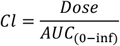

*VD* = *MRT*_(0-inf)_ × *CL* Where,

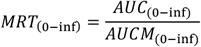

Then, we used the allometric scaling log (VD) vs. log (Weight) and log (CL) vs. log (Weight) to estimate human VD and CL without adjusting for the protein binding. The 3-species fit produced a good correlation with the exponents of 0.77 for VD (R^2^=0.99) and 0.54 (R^2^=0.96) for CL vs. the suggested exponent of <1 for Vd and 0.75 or less for the clearance (**Supplementary Table 1**)^22^. The estimated human CL and VD were used in the one-compartment PK model to plot the plasma concentration vs. time profile (**Figure 3d**). The human MRSD was estimated based on the FDA guidance^23^ and applied to the one-compartment PK model to plot the plasma concentration vs. time profile for multiple SC doses at 6.12 mg/kg QD (**Supplementary Figure S3**).

## Acknowledgment

Following grants were used to support this work: U.S. National Institutes of Health (NIH) grant 1R01CA231509-01A1 to S.W.M., the American Cancer Society Research Scholar Award RSG-15-145-01-CDD to F.L.M, and the Andrew Sabin Family Foundation Fellows Award to F.L.M, the Brockman Medical Research Foundation to F.L.M, the Marine Rose Foundation to FLM, and Uncle Kory Foundation funds to FML supported the work. Y.B. was supported by the CPRIT Training Award (RP210028), Dr. John J. Kopchick Fellowship, and Schissler Foundation. S.K. was supported by the CPRIT Training Award (RP210028) and Larry Deaven Fellowship. K.H. was supported by PCCSM. K.C. and S.K. were supported in part by NIH R01 CA225955 to R. DePinho. We thank Dr. Raghu Kalluri’s lab members for their help and support. We will also thank Dr. Kumaralal K. Kaluarachchi, the manager of the NMR facility at U.T.M.D. Anderson and Dr. Clemens Anklin from Bruker BioSpin. We also acknowledge Dr. Dimitra K. Georgiou and Kenisha Arthur for their help. Illustrations are created with BioRender.com.

## Author Contributions

Y.B. performed NMR, PK studies and generated figures. Y.B., S.K., J.D. and, K.C. assisted with animal works. V.C.Y., M.N.U., and C.D.P. assisted with drug synthesis. K.H. helped with tissue extraction. R. Kalluri and R. Avritcher intellectual inputs. E. J. E. performed pharmacokinetics analysis. S.W.M. assisted with manuscript preparation and data analysis. Y.B. and F.L.M. analyzed data, conceived and wrote the manuscript.

## Supplementary Information

**Supplementary Figure S1:**
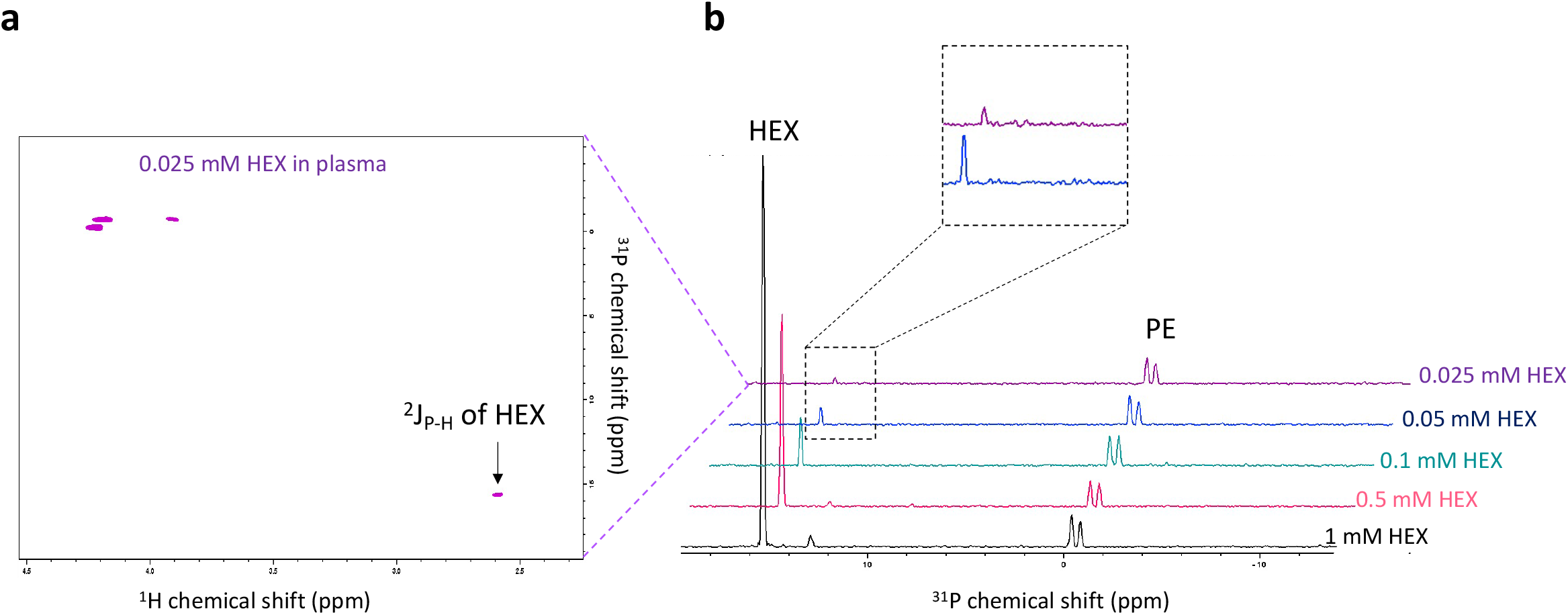
Known concentration of HEX spiked in non-treated NHP plasma to determine the limit of detection and dose calibration curve. Two hundred microliter of plasma were spiked with the known concentration of HEX (0.025 μM to 1 mM) and extracted and prepared for the NMR studies as described in the method section. (**a**) The ^1^H-^31^P HSQC spectrum of the extract of 200 μL of plasma spiked with 50 μM HEX. (**b**) The positive projection of all columns of ^1^H-^31^P HSQC spectra of plasma spiked with 1 mM, 0.5 mM, 0.1 mM, 50 μM, and 25 μM of HEX. The peak integrals of the ^31^P projection were used to calculate the dose calibration curve shown in **Figure 2a**. The detection limit was determined to be 25 μM (for 200 μL plasma, HSQCETGP pulse program, ns=128, and regular 5 mm NMR tube).

**Supplementary Figure S2:**
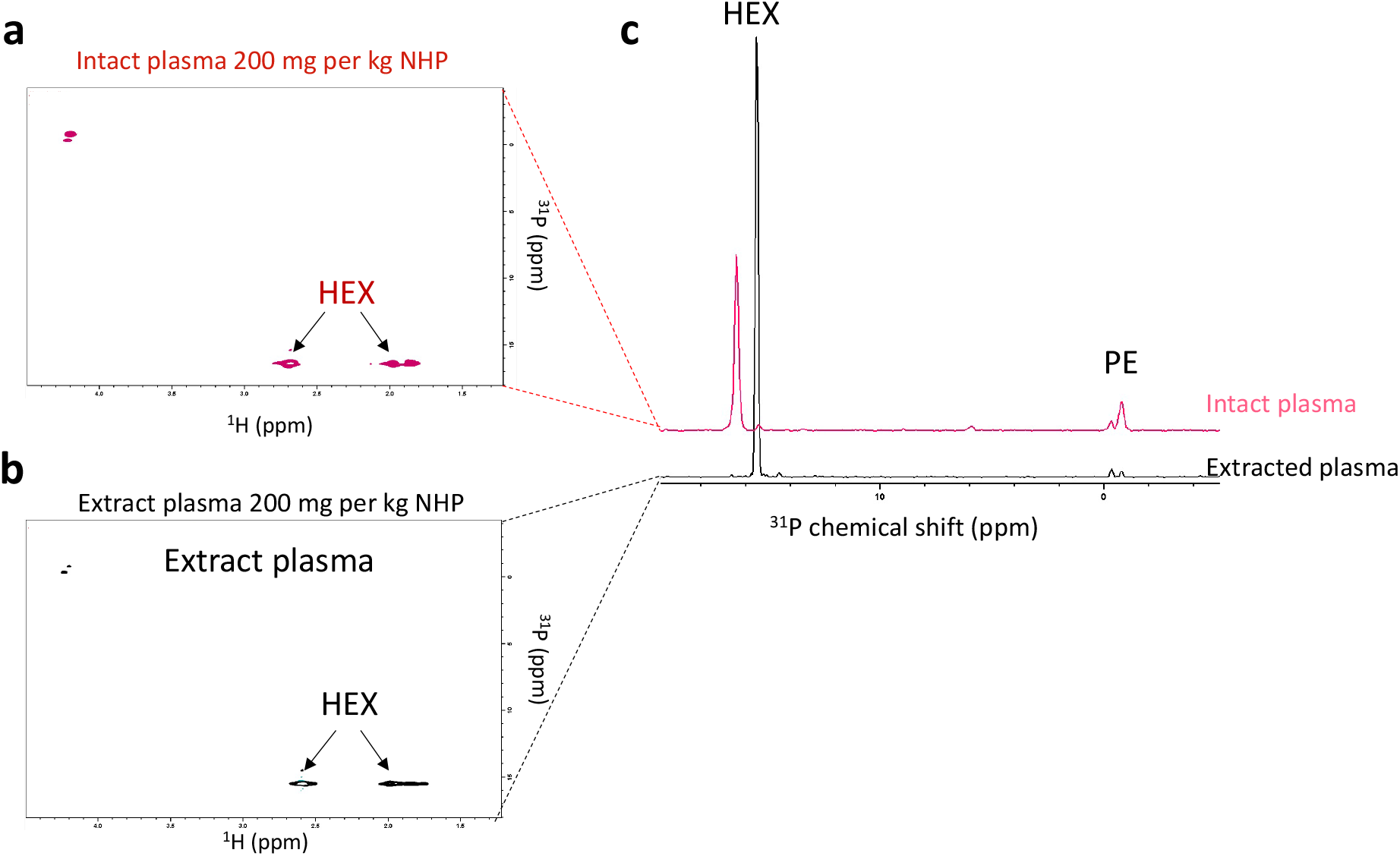
HEX peaks can be detected in the ^1^H-^31^P HSQC spectrum of intact plasma. An NHP was injected with 200 mg per kg of HEX, and blood was drawn one-hour post-injection. The ^1^H-^31^P HSQC spectra of (**a**) 200 μL intact plasma and (**b**) 200 μL of extracted plasma. (**c**) The positive projection of all columns of ^1^H-^31^P HSQC spectra in **a** and **b**. The difference in the microenvironment, PH, and plasma protein binding between the not extracted and extracted plasma result in lower HEX peaks intensity and the change in ^31^P chemical shift observed in intact plasma sample.

**Supplementary Figure S3:**
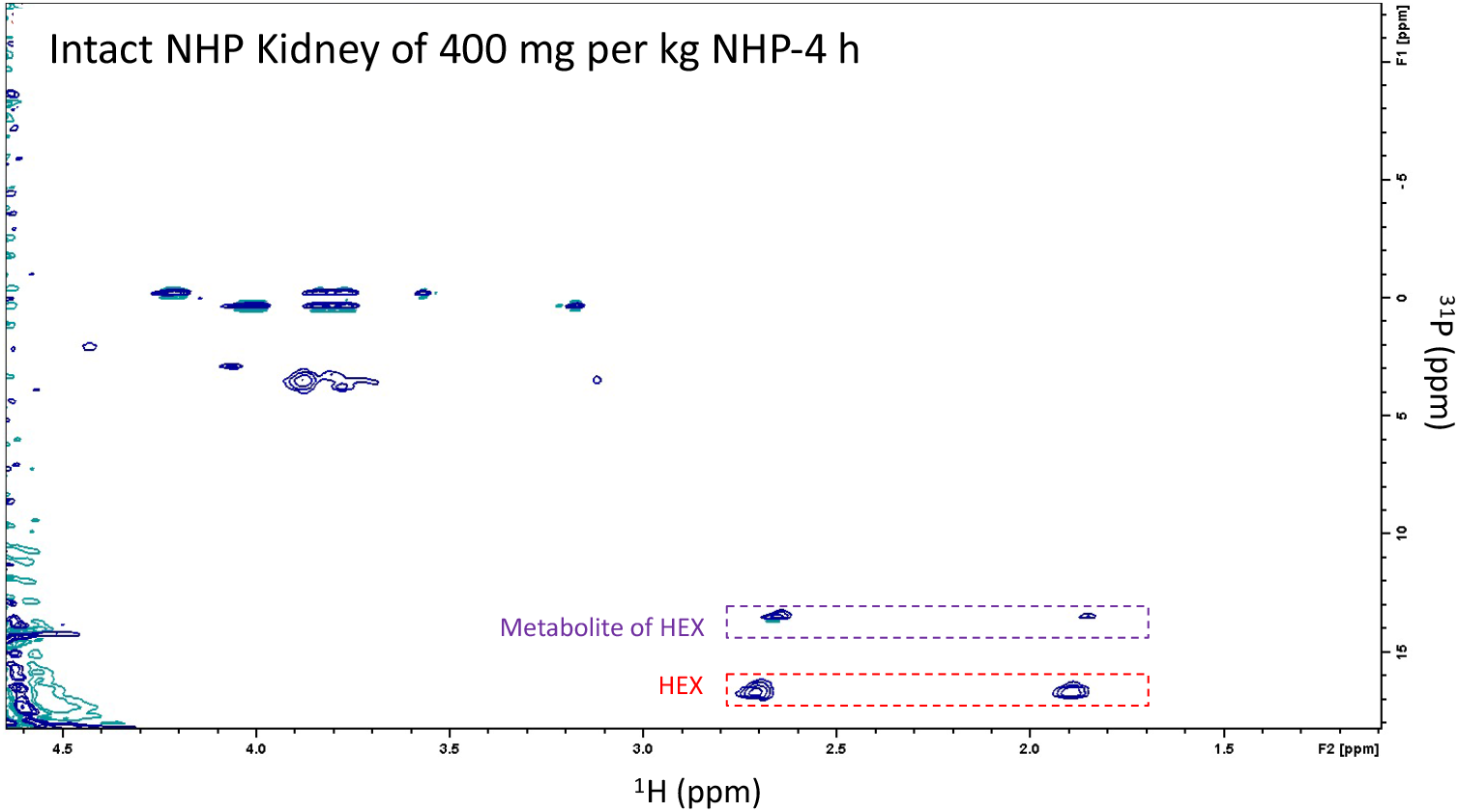
HEX can be detected in the ^1^H-^31^P HSQC spectrum of the intact kidney. An NHP was injected with 400 mg per kg of HEX and euthanized 4 hours later. Two hundred mg of kidney was cut and placed in an NMR tube. The ^1^H-^31^P HSQC spectrum acquired using HSQCETGPSI pulse sequence with ns=128. The spectrum of the extracted sample using the same parameters is shown in **Figure 2g**. This figure suggests that the ^1^H-^31^P HSQC method described in this paper can be applied *in vivo* to study the drug distribution.

**Supplementary Table 1:** Human clearance (CL) and volume of distribution (VD) estimated by allometric scaling. The estimated human PK parameters (CL and VD) were used in the one-compartment PK model to plot the plasma concentration vs. time profile in humans based on a single 150 mg per kg subcutaneous dose. The estimated profile is plotted in **Figure 3d**.

| Parameter | Estimated Human PK Parameters |  |
| --- | --- | --- |
|  | VD (L/kg) | CL (L/h/kg) |
| Value | 0.207 | 0.040 |
| Exponent | 0.77 | 0.54 |
| R<sup>2</sup> | 0.99 | 0.96 |

**Supplementary Table 2.** Conversion of animal doses to HED (human equivalent dose) and MRSD (maximum recommended starting dose) based on BSA (body surface area).

| Species | Weight (kg) | BSA (m<sup>2</sup>) | $K_m$ factor | Equivalent Dose (mg/kg) | HED (mg) | MRSD (mg/kg) |
| --- | --- | --- | --- | --- | --- | --- |
| Cyno Monkey | 2.75 | 0.24 | 11.46 | 1.00 | 200 |  |
| Human Adult | 60 | 1.6 | 37.50 | 0.31 | 61.2 | 6.12 |

**Supplementary Figure S4:**
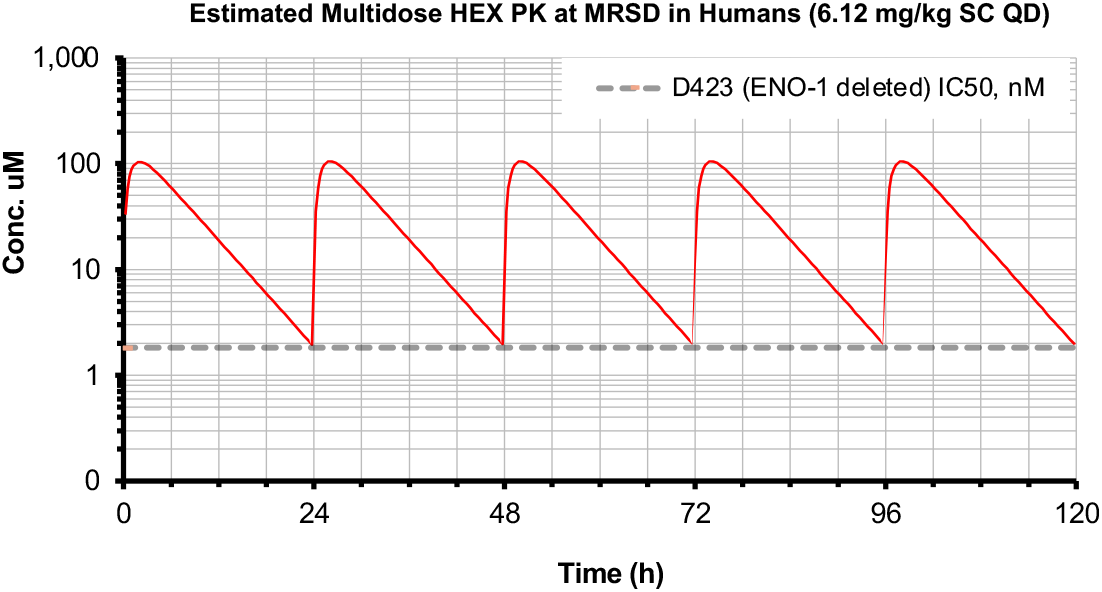
Estimated multidose HEX PK at MRSD in humans (6.12 mg/kg SC QD). The human MRSD (**Supplementary Table 2**) was applied to the one-compartment PK model to plot the plasma concentration vs. time profile for multiple SC doses at 6.12 mg/kg QD.

